# Deep learning models predict regulatory variants in pancreatic islets and refine type 2 diabetes association signals

**DOI:** 10.1101/760868

**Authors:** Agata Wesolowska-Andersen, Grace Zhuo Yu, Vibe Nylander, Fernando Abaitua, Matthias Thurner, Jason Torres, Anubha Mahajan, Anna L Gloyn, Mark I McCarthy

**Affiliations:** Wellcome Centre for Human Genetics; Oxford Centre for Diabetes, Endocrinology and Metabolism; University of Oxford, Oxford, UK; Oxford NIHR Biomedical Centre, Churchill Hospital, Oxford, UK

## Abstract

Genome-wide association analyses have uncovered multiple genomic regions associated with T2D, but identification of the causal variants at these remains a challenge. There is growing interest in the potential of deep learning models - which predict epigenome features from DNA sequence - to support inference concerning the regulatory effects of disease-associated variants. Here, we evaluate the advantages of training convolutional neural network (CNN) models on a broad set of epigenomic features collected in a single disease-relevant tissue – pancreatic islets in the case of type 2 diabetes (T2D) - as opposed to models trained on multiple human tissues. We report convergence of CNN-based metrics of regulatory function with conventional approaches to variant prioritization – genetic fine-mapping and regulatory annotation enrichment. We demonstrate that CNN-based analyses can refine association signals at T2D-associated loci and provide experimental validation for one such signal. We anticipate that these approaches will become routine in downstream analyses of GWAS.

## Introduction

Genome-wide association studies (GWAS) have identified over 400 independent signals implicated in genetic susceptibility to type 2 diabetes (T2D)(Mahajan et al., 2018). However, efforts to derive biological insights from these signals face the challenge of identifying the functional, causal variants driving these associations within the sets of credible variants defined by the linkage disequilibrium (LD) structure at each locus. It remains far from trivial to assign mechanisms of action at these loci: most associated variants map to non-coding sequence, the implication being that they influence disease risk through the transcriptional regulation of one or more of the nearby genes.

There is mounting evidence that disease-associated variants are likely to perturb genes and regulatory modules that are of specific importance within disease-relevant cell types or tissues (GTex Consortium et al., 2017; Marbach et al., 2016). For example, several studies have reported significant enrichment of T2D GWAS variants within pancreatic islet enhancer regions (Parker et al., 2013; Pasquali et al., 2014), with that enrichment particularly concentrated in subsets of islet enhancers characterised by open chromatin and hypomethylation (Thurner et al., 2018), and clustered in 3D enhancer hub structures (Miguel-Escalada, 2018). Pancreatic islets represent a key tissue for the maintenance of normal glucose homeostasis, and uncovering islet-specific regulatory mechanisms is therefore critical to understanding T2D aetiology and pathogenesis.

Several studies have generated genome-wide epigenomic profiling datasets of whole human pancreatic islets, and/or FACS-sorted individual islet cell types (Ackermann, Wang, Schug, Naji, & Kaestner, 2016; Bhandare et al., 2010; Bramswig et al., 2013; Gaulton et al., 2010; Maher, 2012; Parker et al., 2013; Pasquali et al., 2014; Stitzel et al., 2010; Thurner et al., 2018). This wealth of genomic data provides a valuable resource for studying the regulatory machinery of human pancreatic islets, and has proven instrumental in prioritizing disease-associated variants by considering their overlap with regulatory elements enriched in disease-associate signals (Huang et al., 2017; Thurner et al., 2018). However, in cases where multiple associated variants in high LD reside in the same regulatory region, a method with higher resolution is needed to resolve the causal variant. High-throughput massively parallel reporter assays (MPRA) offer one solution for the empirical assessment of putatively functional variants (Tewhey et al., 2016; Ulirsch et al., 2016), but they are expensive to deploy genome-wide, and may not fully recapitulate the cellular context.

Convolutional neural networks (CNNs) are emerging as a powerful tool to study regulatory motifs in genomic data, and are well suited to extracting high-level information from high-throughput datasets *de novo*. Indeed, CNN frameworks have been shown to aid in prioritization of genomic variants based on their predicted effect on chromatin accessibility and modifications (Kelley, Snoek, & Rinn, 2016; Zhou & Troyanskaya, 2015) or gene expression (Kelley et al., 2018; Zhou et al., 2018). The methods deployed so far learn the regulatory code *de novo* from genomic sequences of regulatory regions gathered from multiple tissues, using datasets provided by the ENCODE (Maher, 2012), the NIH Epigenome Roadmap (Bernstein et al., 2010) and GTEx (GTex Consortium et al., 2017) consortia. The derived models offer computational predictions of the likely regulatory effects of genomic variation based on disruption or creation of regulatory motifs discovered by the CNNs. While these multi-tissue methods offer an attractive, generally applicable, framework for variant prioritization, they may be missing nuances of the tissue-specific regulatory grammar, and may not be optimal for predictions of regulatory effects that are specific to disease-relevant tissues.

In the present study, we trained CNNs on a broad collection of genome-wide epigenomic profiles capturing chromatin regulatory features from human pancreatic islets, and applied the resulting models to predict the regulatory effects of sequence variants associated with T2D. We demonstrate that these tissue-specific CNN models recapitulate regulatory grammar specific to pancreatic islets, as opposed to discovering regulatory motifs common across multiple tissues. We apply the CNN models to predict islet regulatory variants among the credible sets of T2D-association signals, and demonstrate how CNN predictions can be integrated with genetic and functional fine-mapping approaches to provide single-base resolution of functional impact at T2D-associated loci.

## Results

### CNNs achieve high performance in predicting islet chromatin regulatory features

We collected 30 genome-wide epigenomic profiling annotations from human pancreatic islets, and their FACS-sorted cell subsets from previously published studies (Ackermann et al., 2016; Bhandare et al., 2010; Bramswig et al., 2013; Gaulton et al., 2010; Maher, 2012; Parker et al., 2013; Pasquali et al., 2014; Stitzel et al., 2010; Thurner et al., 2018)(STable 1), and re-processed them uniformly with the same computational pipelines. The 1000bp long genomic sequences encompassing the signal peaks were used, together with vectors representing presence/absence of the 30 islet epigenomic features within these regions, as inputs to train the multi-class prediction CNNs. The resulting CNN models predict presence of these 30 features within any 1000bp long genomic sequence. Since the weights in neural network training are initialized randomly and then optimized during training, there is a considerable amount of heterogeneity in the predictive scores resulting from different iterations of the same training process, as networks may converge at different local optimal solutions. To improve robustness of results achieved with these models, we trained a total of 1000 CNNs with 10 different sets of hyperparameters differing in numbers of convolutional filters and their sizes to account for this stochastic heterogeneity (STable 2).

Overall, CNNs achieved high performance in predicting the islet epigenomic features in sequences withheld from training, though we observed that the performance varied depending on the predicted feature (Fig.1). The best predictive performance was achieved for features related to promoters, transcription factor (TF) binding and DNA accessibility with mean areas under receiver-operator curves (AUROC) of 0.948, 0.887 and 0.876, respectively. Histone mark features associated with active, coding regions, repressed regions and enhancers proved more difficult to predict based on their underlying genomic sequence, with mean AUROCs of 0.835, 0.792 and 0.777, respectively. As predicted classes were not well balanced by design, we further investigated predictive performance by inspecting the area under precision-recall curves (AUPRC) (Figure 1–figure supplement 1), a more appropriate measure where there are large numbers of negative examples. We observed that, while variable between features, the predictive performance of the CNN models for the chromatin regulatory features was high, and even for the least confidently predicted features far exceeded the performance of random predictors.

**Figure 1.**
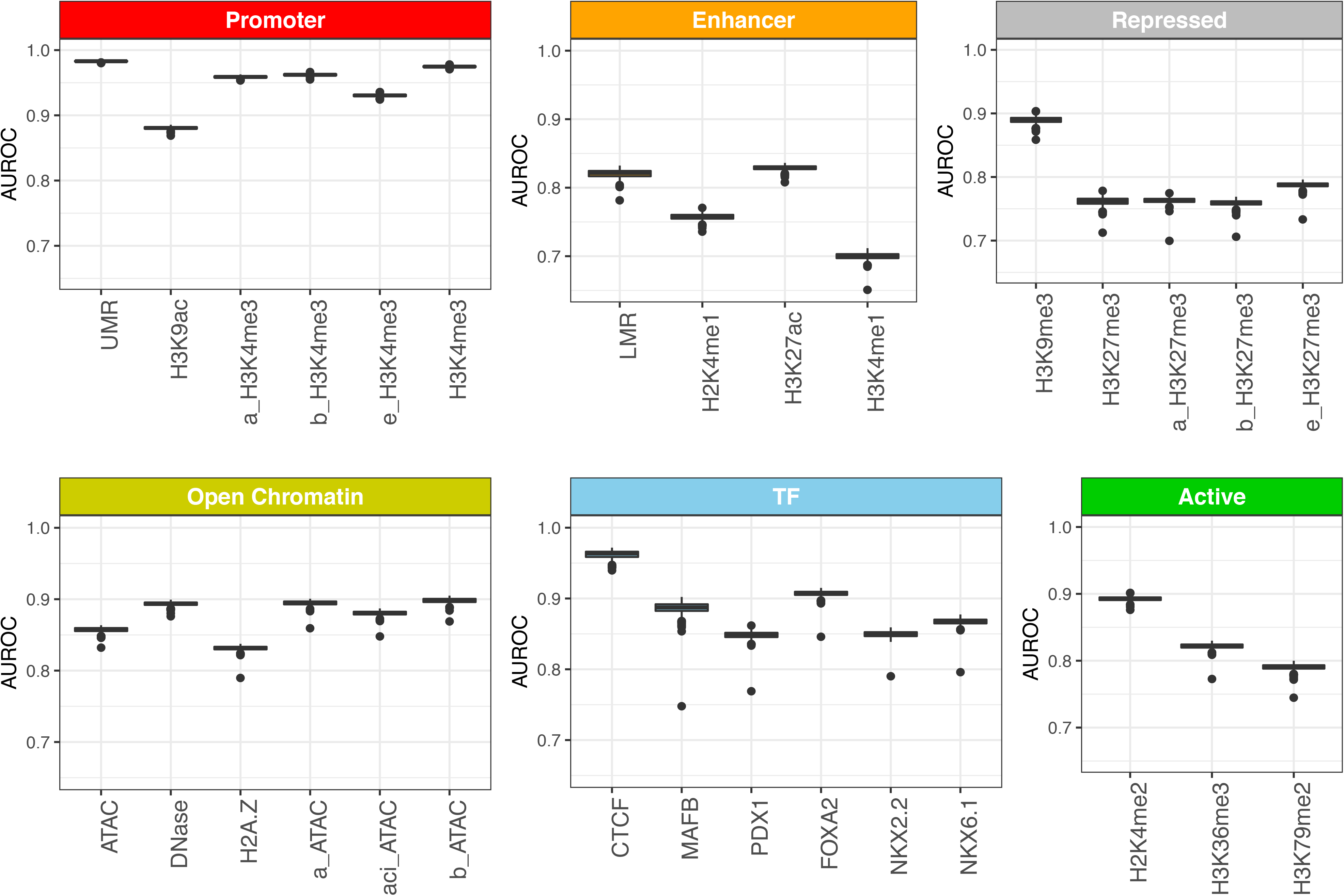
Area under receiver-operator curves (AUROC) for 30 islet epigenomic features predicted by CNN models. The AUROC values were calculated based on performance on the validation set formed by 1000bp sequences from chr2, held out from training and testing. The boxplots show summary of performance across 1000 individual CNN models, and are grouped by corresponding regulatory element.

### Convolutional filters recover binding motifs of TFs with roles in pancreatic development

Convolutional filters of the first network layer capture local sequence patterns and motifs, aiding in predictions of islet regulatory features. We hypothesized that many of these would correspond to binding motifs of transcription factors (TFs) with roles in pancreatic islet development and function. For each convolutional filter of the first CNN layer, we derived a position weighted matrix (PWM) based on the observed nucleotide frequencies activating the filter, and compared them to a database of known TF binding motifs. We observed that the number of annotated motifs per network was positively correlated with the size of convolutional filters within the first layer, while the number of filters informative for the predictions (activation standard deviation>0) decreased with filter size (Figure 1–figure supplement 2). On average, only 29 out of 320 filters in first CNN layer were annotated to known TF binding motifs, but an average of 177 filters were informative for predictions, indicating that CNN models identify potentially novel sequence motifs not currently represented in the databases of known TF binding motifs.

In total, we identified 373 recurrent annotated binding motifs with <5% false-discovery rate (FDR) sequence similarity to the filters of the first CNN layer, which were detected in >50 networks (STable 3). The 10 most frequently discovered non-redundant motifs are listed in Table 1. As expected, among the consistently-detected transcription factor motifs, we found motifs for all the transcription factors included in the ChIP-seq training datasets (CTCF, FOXA2, PDX1, MAFB, NKX2.2, NKX6.1). Additionally, CNNs discovered, *ab initio*, binding motifs for several TFs that are known to be important for pancreatic development and for the maintenance of beta and alpha cell functions, including RFX6, HNF1A and NEUROD1 (Jennings, Berry, Strutt, Gerrard, & Hanley, 2015; van der Meulen & Huising, 2015). This demonstrates that the CNN models of the pancreatic islet epigenome are capable of discovering well-established islet TF motifs *ab initio* from genomic sequences.

**Table 1.**
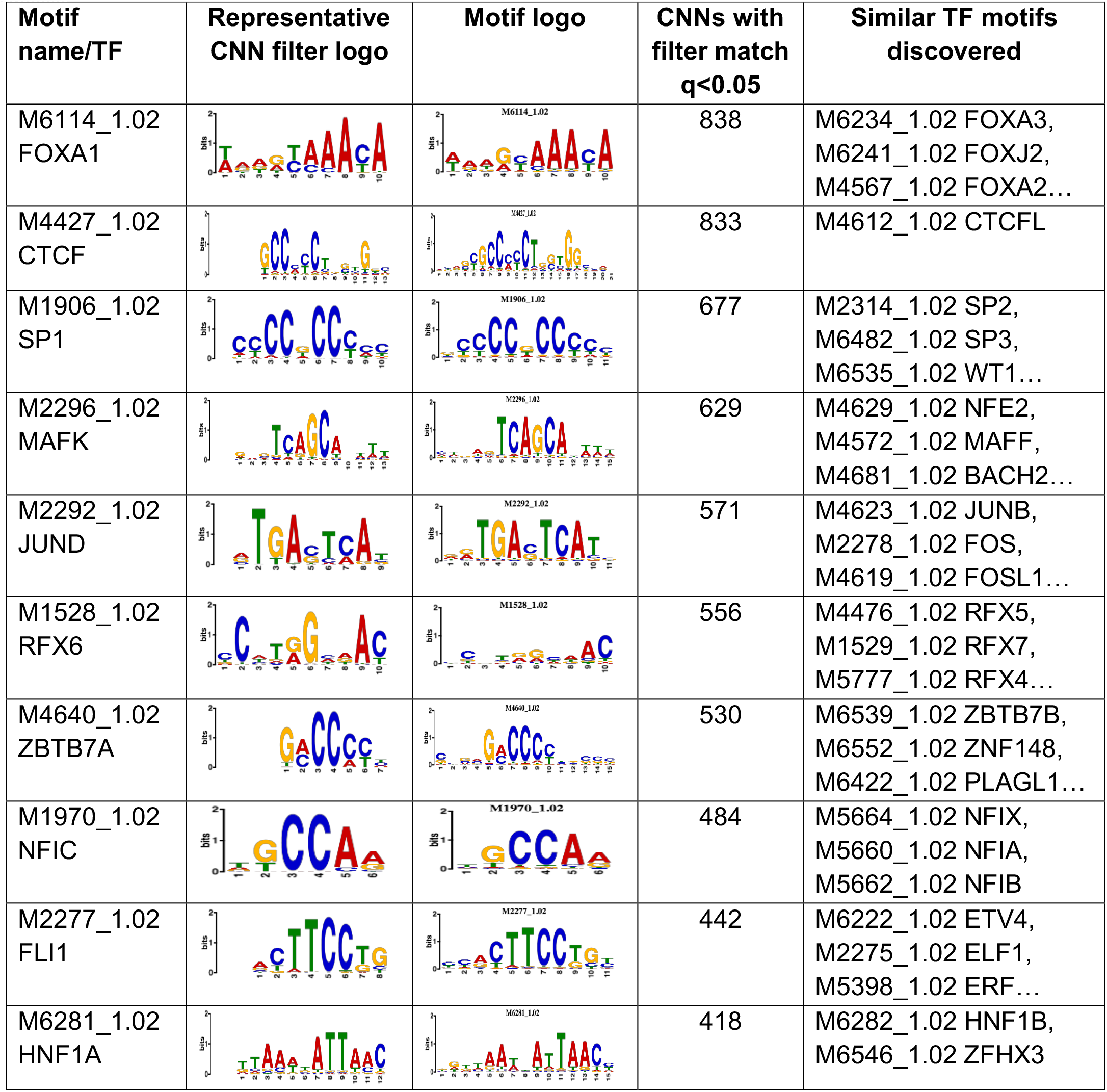
10 non-redundant transcription factors binding motifs most frequently detected by first layer convolutional filters at FDR<5%. Sequence logos of representative CNN filters are shown. Transcription factor binding motifs redundancy was removed with Tomtom motif similarity search with other motifs detected by CNNs with q<0.05 for similarity to the main motif are listed in the last column; only 3 motifs with highest similarity are listed.

### Islet CNN models prioritize T2D-associated variants with regulatory roles in pancreatic islets

The regulatory effects of genomic variants can be approximated by comparing the CNN predictions for genomic sequences including different alleles of the same variant. Here, we applied the islet CNN models for prioritization of T2D-associated variants from a recently published GWAS study (Mahajan et al., 2018). This study of ∼900,000 cases and controls of European ancestry, identified 403 T2D-risk signals, and performed genetic fine-mapping for 380 of them. We ran CNN predictions for all 109,779 variants included within the 99% credible variant sets for these signals, averaging the regulatory predictions for each variant and feature across 1000 individual CNN models to increase robustness. Variants most likely to influence the islet epigenome were then identified through the cumulative distribution function for the normal distribution, separately for each predicted feature, and the lowest q-value for any of the features was assigned to a variant to signify its overall regulatory potential.

We identified a total of 11,389 variants with a q-value<0.05 for any of the 30 features, approximately 10% of the total number of credible set variants. These variants were significantly more likely to be evolutionary conserved than the credible set variants with overall q>0.05, as assessed by one-sided Wilcoxon rank sum test of the GERP scores (Cooper et al., 2005) (p=7.31e-04).

We reasoned that variants predicted to affect function of specific regulatory elements would be more likely to reside within them, e.g. a variant predicted to disrupt enhancer function should be residing in an enhancer region. We tested this by investigating overlap of predicted regulatory variants with human pancreatic islet chromatin state maps (Thurner et al., 2018). For this purpose, we considered the lowest q-values within each of the earlier described groups of regulatory features, corresponding to promoters, enhancers, open chromatin, transcription factor binding, as well as active and repressed regulatory regions (Fig.2). Overall, we observed good agreement between the predicted disrupted regulatory elements and the variant located within them. Additionally, we observed a depletion of regulatory variants within heterochromatin and other low methylation sites.

**Figure 2.**
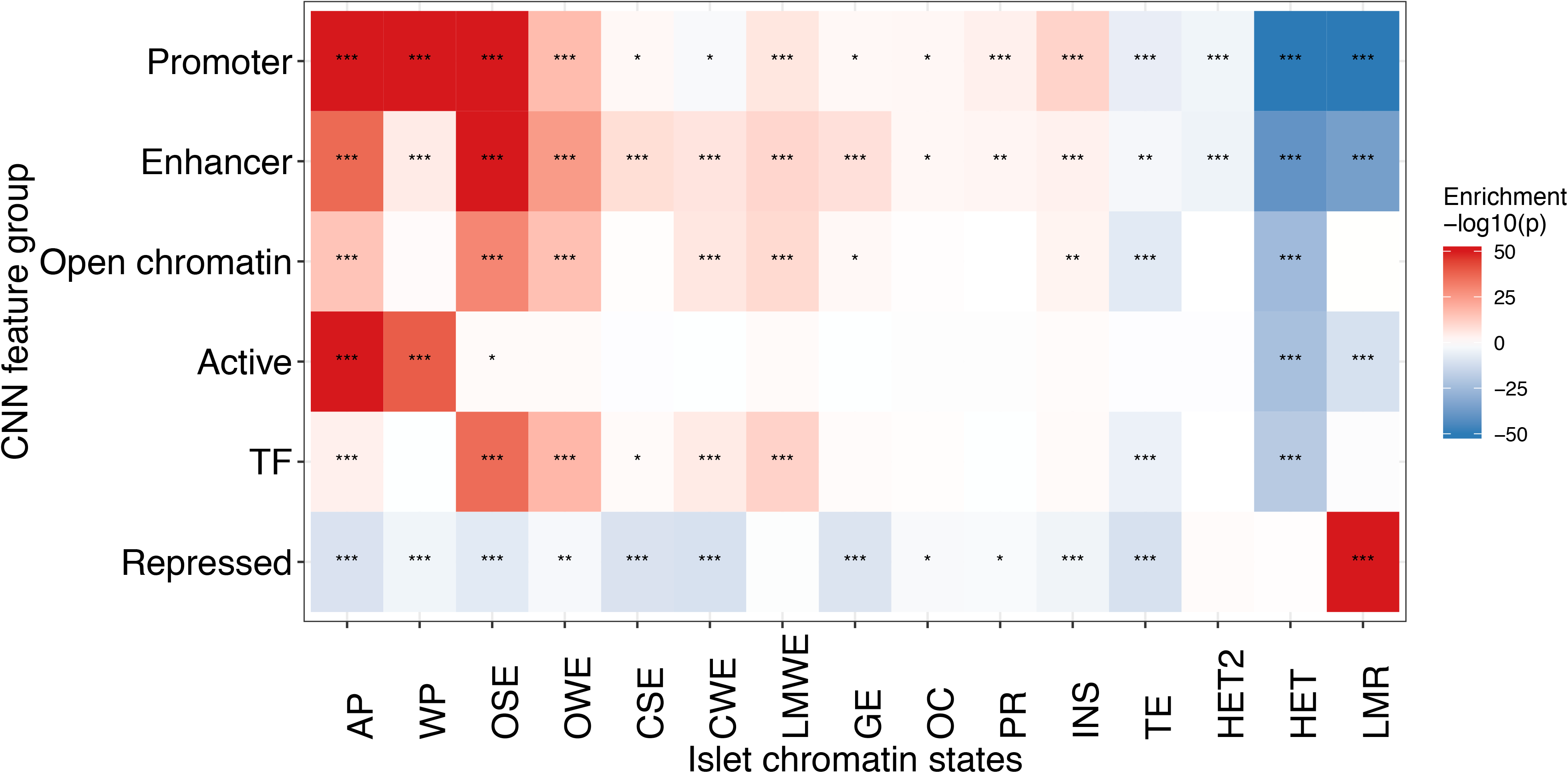
Predicted regulatory variants reside in regulatory elements they are predicted to affect. For each variant we found the lowest CNN q-value among feature groups corresponding to different regulatory elements (promoters, enhancers, open chromatin, active regions, TF binding, repressed regions) predicted from genomic sequence, and we ranked all variants according to these six q-values. We then tested whether variants residing in each of the 15 pancreatic islet chromatin states (Thurner et al., 2018) were enriched at the top or bottom of these ranked lists. Colours in the heatmap represent the strength of the enrichment expressed as log10-transformed enrichment q-values, with red colours representing enrichments at the top (enrichment), and blue at the bottom of the ranked lists (depletion). For plotting purposes all −log10(p-values) below −50, or above 50 were truncated to these values. Stars denote significant enrichments: * <0.05, ** <0.01 and *** <0.001.

Finally, we hypothesized that predicted regulatory variants would be more likely to show allelic imbalance in chromatin accessibility. We used ATAC-seq data from a previously-published dataset of 17 human pancreatic islets (Thurner et al., 2018) to identify 137 pancreatic islet chromatin accessibility QTLs (caQTLs) among the credible set variants, and found these to be significantly enriched among the variants with the lowest CNN q-scores (p=6.93e-20).

### Convergence between CNN predictions and fine-mapping approaches

If the CNN models are correctly identifying regulatory variants, we would expect to see convergence between variants predicted to have regulatory effects based on the CNNs, and those assigned high genetic posterior probabilities of association (gPPA) from genetic fine-mapping. To test for such convergence, we generated the null distribution of randomly distributed regulatory variants through 1000 permutations of CNN q-values, while preserving the structure of credible sets at the 380 T2D-associated signals. We observed enrichment of islet regulatory variants (CNN q<0.05) among variants with highest PPAs (Fig.3A), compared to the permutation-based random distribution of regulatory variants (p=0.001). Overall, we found that 28.8% of variants with gPPAs>=0.8 had predicted regulatory effects with q<0.05.

**Figure 3.**
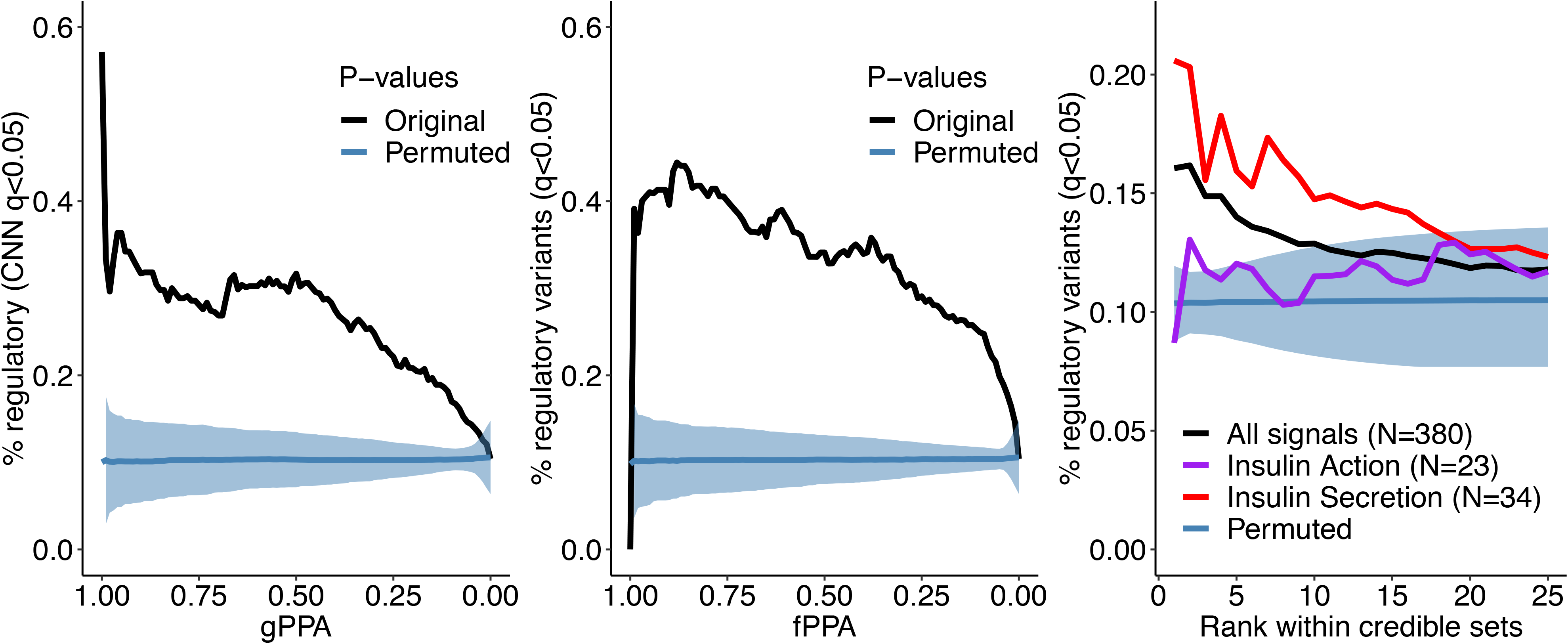
Convergence between CNN regulatory predictions and fine-mapping approaches for functional variant prioritization. A. Regulatory variants (black) are enriched among variants with highest genetic PPAs (gPPAs) over permuted background (blue). B. Regulatory variants (black) are enriched among variants with highest functional PPAs (fPPAs) generated with FGWAS over permuted background (blue). C. Regulatory variants (black) are enriched among variants with top PPA ranks within sets of credible variants over permuted background (blue), as well as at top ranks of signals acting through insulin secretion (red) over insulin action (purple) mechanisms.

It is standard to complement genetic fine-mapping with information from the genome-wide enrichment of association signals within regulatory annotations in disease-relevant tissues, deriving functional posterior probabilities of association (fPPAs) that combine genetic and epigenomic insights into variant function (Huang et al., 2017). As with the gPPAs, we observed that variants with high fPPAs, obtained through incorporating enrichments in regulatory elements from human pancreatic islet chromatin state maps (Thurner et al., 2018), were enriched for CNN-predicted regulatory effects (p=0.001)(Fig.3B). Overall, 40.6% of variants with fPPAs>=0.8 had predicted regulatory effects with q<0.05.

As fine-mapping resolution varies between signals, we conducted analogous analyses based on variant rank within each credible set, irrespective of the quantitative PPA value (Fig.3C). Again, we observed higher proportions of predicted regulatory variants at higher PPA ranks within fine-mapped loci, when compared to random distribution of regulatory variants.

One might expect that CNN models trained with pancreatic islet epigenomic annotations would display the strongest evidence for prediction of regulatory variants at the subset of T2D GWAS signals characterized by defects in insulin secretion (Dimas et al., 2014; Wood et al., 2017), signals that are likely mediated through events in pancreatic islets. Indeed, we observed that the enrichment of islet CNN-regulatory variants was more marked within the top ranks of insulin secretion signals. In contrast, T2D signals characterized by a primary defect in insulin action (which typically involve mechanisms in liver, fat and muscle) showed no enrichment over the permuted background (Fig.3C).

Collectively, these data corroborate the convergence of the agnostically-derived CNN variant regulatory scores and diverse measures of islet biology, and indicate the potential for CNNs to support causal variant prioritization in T2D. They also emphasize the value of functional analyses that take account of the tissue-specificity of both transcriptional regulation and disease pathogenesis.

### Islet CNN models correctly identify known functional variants at T2D-associated loci

We tested whether the CNN regulatory predictions for individual variants can be integrated with previous fine-mapping approaches to further resolve T2D-associated signals. Overall, we found that, at 327 out of the 380 fine-mapped signals, there was at least one predicted regulatory variant (defined as q<0.05). Among the 74 signals previously fine-mapped to a single variant (with gPPA or fPPA>=80%), we found 28 variants predicted to be regulatory in pancreatic islets (STable 4), in line with the previously reported overall enrichment of T2D-association signals in the regulatory elements specific to islets (Miguel-Escalada, 2018; Parker et al., 2013; Pasquali et al., 2014; Thurner et al., 2018). These included two well-studied variants (rs10830963 at *MNTR1B*, rs7903146 at *TCF7L2*), both previously shown to alter enhancer activities (Gaulton et al., 2015; Gaulton et al., 2010): these served as positive controls for the application of CNNs to functional variant prioritization.

### Islet CNN models refine regulatory mechanisms at T2D-associated loci

While functional fine-mapping can be invaluable in narrowing down the list of most likely causal variants through investigating overlaps with regulatory elements in appropriate tissues, this strategy may not provide sufficient resolution at loci where several variants reside in the same regulatory element. Among the credible sets of variants, we identified 93 signals featuring at least two variants with fPPAs>=20%, indicating that, even after functional fine-mapping, there is no unambiguously causal single variant. At 37 of these 93 signals, CNNs predicted regulatory variants among the top fPPA candidates (STable 5). At 25 signals, integration of CNN regulatory predictions downstream of the functional fine-mapping highlighted a single most likely causal variant, with either just a single regulatory variant predicted among the credible variants, or the top regulatory variant having much lower q-value (difference in −log10(q) >100) than other predicted regulatory variants at the signal (few such signals are highlighted in Fig.4 and Fig.5), narrowing down the list of candidates for further functional follow-up studies.

**Figure 4.**
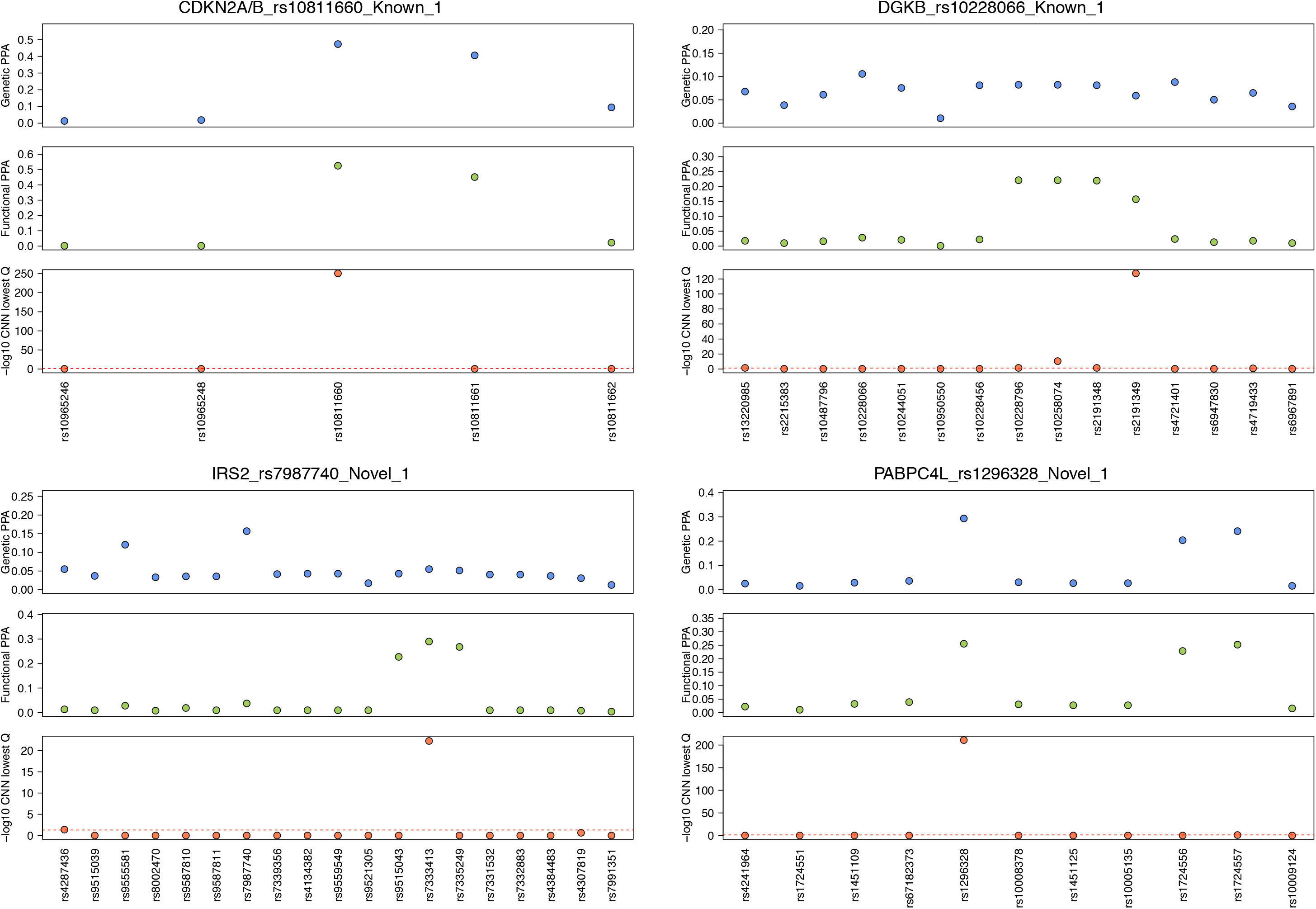
Examples of T2D-association signals where integration of CNN regulatory variant prediction downstream of functional fine-mapping refines the association signals to single candidate variants. Genetic PPAs (gPPAs) are shown in the top panels as blue points, functional PPAs (fPPAs) are shown in the middle panels as green points, and −log10-transformed q-values from CNN predictions are shown in the bottom panels as red points.

**Figure 5.**
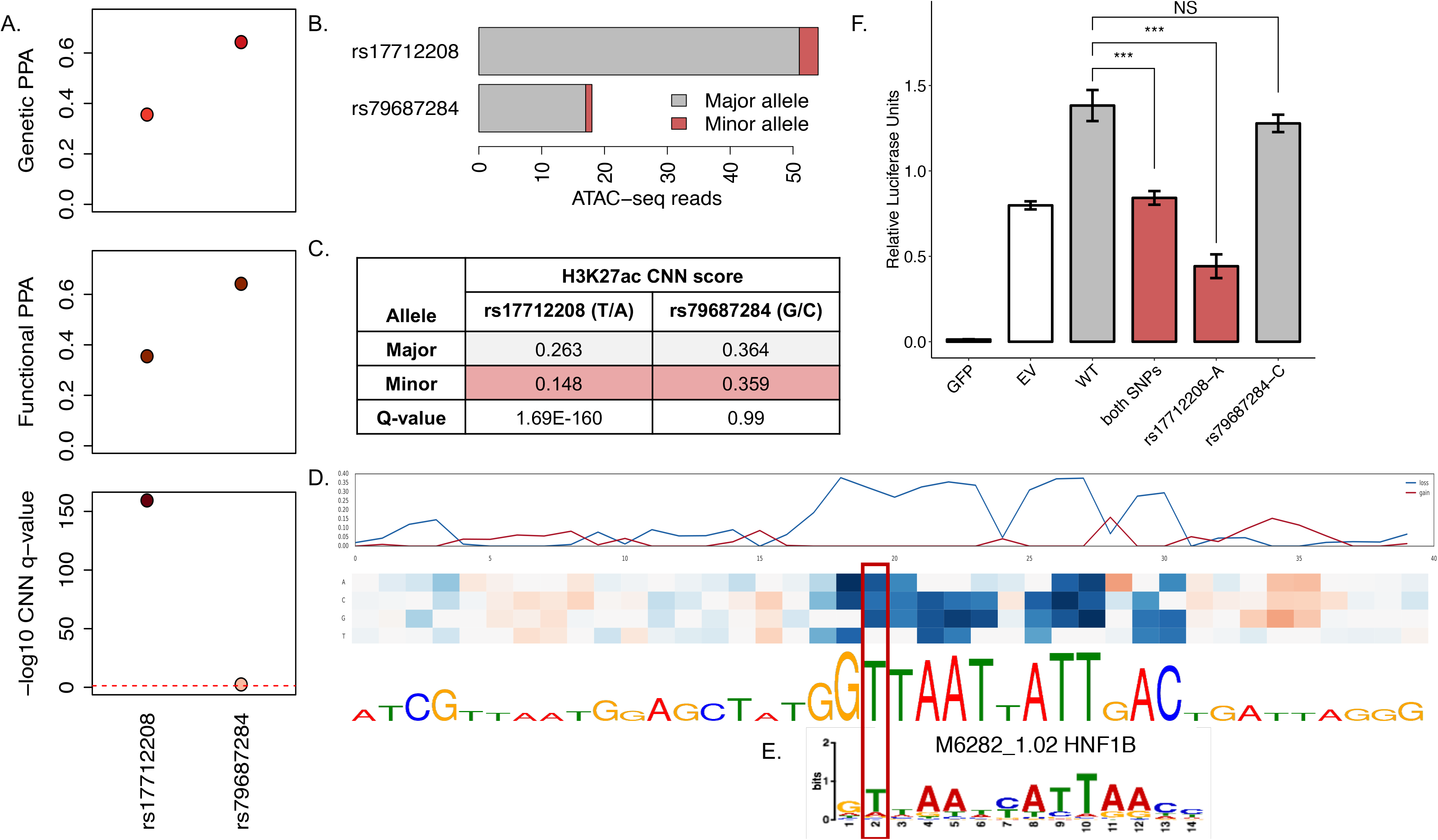
CNN regulatory predictions help refine the association signal at PROX1 locus, previously fine-mapped to only two variants: rs17712208 and rs79687284. A. Genetic PPA (gPPA), functional PPA (fPPA) and −log10(q-value) of the CNN islet regulatory predictions for both variants. B. Allelic imbalance in open chromatin across 4 pancreatic islets heterozygous for the variants. Allele counts for the major (grey) and minor (red) alleles are shown for both variants. C. Table summary with CNN predictions for the H3K27ac mark for both variants. D. *In silico* saturated mutagenesis for 40nt flanking sequence around the rs17712208 variant for the H3K27ac predictions. The line plots in the upper panel indicate the absolute highest values from the heatmap below, with blue line indicating loss of function, and red – gain of function changes. Blue fields in the heatmap indicate that a given nucleotide substitution would result in decrease in prediction values for H3K27ac, while red field indicate increase in the predictions. The height of letters in the sequence below the heatmap indicated the relative importance of each nucleotide in the final predictions. E. Matched HNF1B binding motif is shown below. F. Luciferase reporter assays confirmed that the A allele of rs17712208 resulted in significant repression of enhancer activity, while no effect was observed for the rs79687284 variant. GFP = green fluorescent protein (negative control), EV = empty vector (baseline).

To explore this further, we focused on a T2D-association signal at the *PROX1* locus, identified after conditioning the T2D association on the primary (most significant) association signal at rs340874, which acts through the insulin secretion pathway. This conditional signal has been fine-mapped to two plausible variants, rs79687284 and rs17712208, on the basis of genetic and genomic data (Fig.5A). These variants are in perfect LD (R^2^=1.0, D’=1.0) and are located 376bp apart within the same open strong enhancer in islets. Neither genetic fine-mapping (in European populations) nor functional fine-mapping were able to further resolve the association at this signal. When we investigated pancreatic islet ATAC-seq data from 4 individuals heterozygous for these two variants (as confirmed by array genotype data), we observed strong allelic imbalance at both variants (rs17712208 p=1.55e-06; rs79687284 p=6.10e-05) (Fig.5B). This supports regulatory effects of this *PROX1* signal in the pancreatic islets, but highlights the difficulties in resolving the causal variant. These data are also consistent with the possibility that both variants are contributing to the functional effect.

At this signal, both variants were scored as potentially regulatory by CNNs with q<0.05, but the q-value at rs17712208 was much lower (q=1.69e-160) highlighting this variant as the more likely regulatory candidate in the islets. This q-value corresponded to a prediction that the T2D risk A-allele would lead to a significant reduction in the H3K27ac mark (△H3K27ac=-0.12) (Fig.5C) indicative of an active regulatory element presence. The second variant, rs79687284, was assigned a much less remarkable q-value (q=0.002) and only predicted to affect promoter features, with the top predictions pointing to a mild reduction of H3K4me3 mark in alpha and beta cells (△a_H3K4me3=-0.01, △b_H3K4me3=-0.01).

While it remains plausible that rs79687284 plays a regulatory function in a specific cellular context, the functional annotation and CNN predictions point to rs17712208 as the more likely causal variant (particularly given that both map to an enhancer rather than a promoter). We performed an *in silico* saturated mutagenesis for the rs17712208 variant (Fig.5D) to identify the regulatory sequence motifs at this locus affected by the variant. We observed that the reference T-allele is a crucial nucleotide in the HNF1B binding motif (Fig.5E), and that introduction of an A-allele at this position disrupts this motif, leading to predicted loss of the H3K27ac mark. Using an *in vitro* reporter assay in the human EndoC-βh1 beta-cell model, we confirmed that the A-allele at rs17712208 resulted in significant repression of enhancer activity (p=0.0001), while no significant change was observed for the rs79687284 variant (p=0.15)(Fig.5F).

These examples illustrate how diverse approaches to variant prioritization at the T2D-associated loci show strong convergence towards the same candidate variants, and how CNN models can complement fine-mapping approaches in providing single-base resolution to refine the association signals. Coupled with additional evidence that these signals exert their mechanism of action in the pancreatic islets, rather than other tissues implicated in T2D aetiology, the predicted regulatory variants may provide attractive targets for further functional follow-up studies.

### Comparison of tissue-specific with multi-tissue CNN model

Finally, we compared the regulatory variant predictions made with the pancreatic islet tissue-specific CNN models described in this study with predictions made with a widely-used multi-tissue variant prioritization tool, DeepSEA (Zhou & Troyanskaya, 2015) (Figure 3–figure supplement 1). We compared our results to the multi-tissue significance scores reported by DeepSEA for each variant, as well as to the predicted effects on pancreatic islet chromatin accessibility (based on ENCODE DNase-seq data [“PanIslets”] generated from primary pancreatic islet cells, one of the tissues contributing to the overall multi-tissue score). Overall, we observed a modest but highly significant correlation of predictions with both these regulatory prediction scores reported by DeepSEA (multi-tissue: r=0.227, p<2.2e-16; PanIslets: r=0.223, p<2.2e-16)(Figure 3–figure supplement 1 - AB).

As with our islet CNN models, we observed strong enrichment of predicted regulatory variants amongst the top ranks of the T2D GWAS credible variant sets for both modes of DeepSEA, as compared to permuted background (Figure 3–figure supplement 1 - C). However, both the multi-tissue and PanIslets DeepSEA models performed better within the subset of T2D signals acting through insulin action rather than through insulin secretion signals, predicting higher proportions of regulatory variants at top ranks of credible sets for the former, contrary to our expectations for the PanIslets model (Figure 3–figure supplement 1 - D). In addition, the DeepSEA framework failed to predict regulatory impact at loci where T2D causative variants have been established through multiple lines of evidence (such as at rs10830962 at *MTNR1B* (Gaulton et al., 2015) and rs7903146 at *TCF7L2* (Gaulton et al., 2010)), both of which were correctly identified as regulatory in our islet-based study. Similarly, while the rs17712208 variant at *PROX1* reported in this study was found to be regulatory using the multi-tissue mode of DeepSEA (p=6.25e-03), the predictions for pancreatic islets accessibility profiles (PanIslets) were only borderline significant for this variant (p=0.049).

Our *in silico* saturated mutagenesis analysis indicated NEUROD1 as the likely motif created at the *MTNR1B* locus (not shown), and HNF1B as the motif disrupted at *PROX1* (Fig.5E). These transcription factors are critical for the function of pancreatic islet beta cells (Gu et al., 2010), and likely represent tissue-specific processes, more liable to be missed in a multi-tissue model. We anticipate that CNN models trained on data from multiple tissues may be biased towards sequence features and motifs present across all the tissues, and may not detect tissue-specific signals with high confidence. Given the demonstrated enrichment of T2D association signals in regulatory elements specific to the pancreatic islets, we demonstrate the value of training the prediction models in the tissues most relevant to the phenotype in question, at least for those loci where the tissue-of-action can be defined.

## Discussion

Application of deep learning methods to characterize the regulatory potential of noncoding variants has been a subject of interest in recent years. Instead of predictions across an array of tissues, in the current study we focused on multiple genome-wide epigenomic annotations available for pancreatic islets, one of the central tissues impacting T2D aetiology. We demonstrate how the CNN regulatory predictions for genomic variants can be integrated downstream of fine-mapping approaches for further refinement of GWAS association signals.

Overall, we observed that CNNs were capable of learning the regulatory code from the underlying genomic sequence, though the prediction accuracy differed among the studied epigenomic features. The individual feature predictive performance was likely affected by a number of factors, in particular the quality of input data, and the total number of peaks called for a feature, as deep learning methods require many training examples to learn to generalize well. Furthermore, epigenomic features differ in their characteristics, including peak width and specificity of sequence motifs predictive of their presence. Thus, when training a multi-task prediction network, we may expect that network parameter optimization might not work equally well for disparate features. Promoter regions, DNA accessibility and binding of specific transcription factors proved the easiest to predict, as these features are characterized by very distinct sequence motifs that can be identified by the network’s convolutional filters. Importantly, the binding motifs learned d*e novo* from the genomics sequences of signal peaks presented to the networks included several transcription factors with well-established functions in pancreatic development and function.

In the present study, we trained 1000 individual CNN models with different hyperparameters to aid in prioritization of the likely functional variants from the largest T2D GWAS study to date (Mahajan et al., 2018). We observed an overall convergence of the genetic fine-mapping, functional fine-mapping and regulatory predictions generated by the tissue-specific CNN models, as evidenced by enrichment of predicted functional variants among variants with highest gPPAs, fPPAs and PPA ranks. To our knowledge, this is the first time such enrichment has been reported for *in silico* predictions of regulatory variants. Our findings contrast with those from a recent study, which argued that computational methods are not yet mature enough to be applied in GWAS fine-mapping (Liu et al., 2019).

In our study, we have observed multiple examples of convergence of the top scoring variants by CNNs with the top variants identified through functional fine-mapping using islet chromatin state maps. We also note that the CNN-prioritized variants were significantly enriched in islet chromatin accessibility QTLs, as well as variants residing in islet functional regulatory elements, including promoters and strong open enhancers, which is in line with findings of previous functional genomics studies (Parker et al., 2013; Pasquali et al., 2014; Thurner et al., 2018). We demonstrate that when used in conjunction with high-resolution genetic or functional fine-mapping, CNN predictions can provide a powerful way of dissecting the causal variant from a set of T2D-associated variants. The results presented in this study will facilitate further functional follow-up studies to fully elucidate the mechanisms underlying the associated disease susceptibility at the prioritized variants. One caveat here is that the value of applying tissue-specific CNN models to guide the identification of causal variants, depends on knowing the effector tissue for the signals in question. With complex diseases, such as T2D, this is not always straightforward, as multiple tissues (including pancreatic islets, liver, adipose and skeletal muscle) have been implicated in T2D aetiology. As more functional data on disease-relevant tissues are collected, and large scale experimental studies provide validated sets of regulatory variants to include in supervised model training, we expect further advancements in the deep learning applications in genomics. Indeed, we anticipate that in the near future, deep learning based fine-mapping approaches will become part of routine downstream analyses of GWAS.

## Materials & Methods

### Collection and processing of islet data

We collected 30 genome-wide epigenome profiling datasets from human pancreatic islets from previously published studies (Bhandare et al., 2010; Gaulton et al., 2010; Maher, 2012; Parker et al., 2013; Pasquali et al., 2014; Stitzel et al., 2010; Thurner et al., 2018)(STable 1). Where raw sequencing data was available, we uniformly reprocessed the data using the default settings of the AQUAS Transcription Factor and Histone ChIP-Seq processing pipeline (https://github.com/kundajelab/chipseq_pipeline) and ATAC-Seq/DNase-Seq pipeline (https://github.com/kundajelab/atac_dnase_pipelines), mapping against the human reference genome build hg19, and using MACS2 for peak calling. We used the coordinates of the optimal peaks sets produced with the Irreproducible Discovery Rate (IDR) procedure for further analysis.

### CNN input sequence and feature encoding

The training and test datasets for the convolutional neural network training were created analogously to the approach used in Basset (Kelley et al., 2016). Briefly, 1000bp genomic intervals were assigned to all called narrow peaks, by extending 500bp on each side of the centre of the peak. Peaks were greedily merged until no peaks overlapped by >200bp, and the centre of the merged peak was determined as a weighted average of the midpoints of the merged peaks, weighted by the number of chromatin features present in each individual peak. This resulted in 505,273 genomic intervals of 1000bp length with assigned presence/absence of the 30 islet epigenomic features, with an average of 2.62 chromatin features per interval. The genomic sequences of the intervals were extracted from the hg19 human reference genome and encoded as one-hot code matrix, mapping the sequences into a 4-row binary matrix corresponding to the four DNA nucleotides at each position. For each 1000bp sequence we created an accompanying feature vector denoting which of the 30 datasets showed a signal peak overlapping the sequence. All sequences from chromosomes 1 (N=43,029, 8.52% total) and 2 (N=40,506, 8.02% total) were held out from the training and applied as test and validation sets, respectively.

### CNN training

CNN models were trained using code from the open source package Basset (Kelley et al., 2016) implemented in the Torch7 framework (http://torch.ch). We trained a total of 1000 CNN models with 10 different hyperparameter settings (Sup. Table 2), differing in sizes and numbers of convolutional filters applied. We averaged the results across these 1000 models to achieve robust results, and overcome the CNN training stochasticity. Each network contained three convolutional layers, each followed by a rectified linear unit (ReLU) and a max pooling layer, and two fully connected layers. The final output layer produced the predictions for the 30 features. The network training was stopped when the area under receiver-operator curve (AUROC) did not improve in 10 subsequent training epochs. The predictive performance of the networks was assessed by evaluating predictions made on the sequences from chr2 constituting the validation set. We calculated the AUROCs and AUPRCs using R package PRROC v.1.3 (Grau, Grosse, & Keilwagen, 2015).

### Sequence motifs captured by convolutional filters

The convolutional filters in the first CNN layer detect repeatedly occurring local sequence patterns, which increase the prediction accuracy. These patterns can be summarized as position-weighted matrices (PWMs), derived by counting nucleotide occurrences that activate the filter to more than half of its maximum value, as implemented in the Basset framework. The resulting PWMs were matched to the CIS-BP database (Weirauch et al., 2014) of known transcription factor binding motifs using the Tomtom v.4.10.1 motif comparison tool (Gupta, Stamatoyannopoulos, Bailey, & Noble, 2007). We repeated this for all the 1000 trained networks, and found recurrent motifs by identifying binding motifs matched at FDR<5% in at least 50 networks. The redundant motifs were filtered out from the top motifs table by comparing all the database motifs against each other using the Tomtom v4.10.1 motif comparison tool, and reporting the top scoring motif as the main hit, and other motifs matching it at FDR<5% as secondary motifs.

### Application of CNN models to variant prioritization

We obtained the credible variant sets from a recent T2D GWAS study (Mahajan et al., 2018). For each variant, we run predictions for the 30 islet epigenomic features with all 1000 trained CNN models for two 1000bp long sequences: first including the reference allele and second including the alternative allele, with the variant positioned in the middle of the 1000bp sequence. We calculated the mean differences in prediction scores across all the 1000 CNNs for each of the investigated features between the two sequences for each variant. We then estimated p-values for each of the variants and each feature with the cumulative distribution function of normal distribution, and applied FDR procedure for multiple testing correction. The overall regulatory potential of each variant was quantified as the lowest q-value for any of the 30 features.

### Functional validation of CNN regulatory predictions

To test the validity of the resulting regulatory predictions for the credible set variants, we tested whether variants predicted to affect specific regulatory elements (promoters, enhancers, open chromatin, TF binding, active, or repressed regions) were more likely to reside in these regulatory regions in islets, as defined by the high-resolution chromatin state maps of human pancreatic islets, partitioning the genome into 15 regulatory states based on patterns of chromatin accessibility and DNA methylation integrated with established ChIP-seq marks (Thurner et al., 2018). Using the generally applicable gene set enrichment R package gage (version 2.26.1)(Luo, Friedman, Shedden, Hankenson, & Woolf, 2009) we tested whether variants overlapping each chromatin state were more likely to be found at the top or bottom of the variant list ranked by the lowest q-values within each of the feature groups.

In a similar manner we tested whether variants with lower q-values are more likely to act as chromatin accessibility QTLs (caQTLs) exhibiting allelic imbalance in open chromatin in the ATAC-seq data from 17 human pancreatic islets (Thurner et al., 2018). We identified the islet caQTLs among the credible set variants by investigating variants overlapping the ATAC-seq peaks. We required >= 2 subjects with a heterozygous genotype for the variant, and >=5 sequencing reads overlapping each allele. We calculated the significance of open chromatin allelic imbalance with the negative binomial distribution test, and used an FDR-adjusted p-value of 0.05 to define the 137 caQTLs. The enrichment of these caQTLs within the credible set variant list ranked by their lowest q-value was calculated using R package ‘gage’.

### Evaluating convergence between fine-mapping and CNN predictions

To test the overall convergence between the two complementary methods for variant prioritization: genetic fine-mapping and CNN regulatory predictions, we evaluated whether we observed more regulatory variants at highest genetic PPAs. We examined the proportions of regulatory variants predicted by CNNs using q-value <0.05 as a cut-off, at different thresholds of genomic PPA (gPPA) going down from PPA of 1.00 to 0.00 in steps of 0.01, and observed decreasing proportions of regulatory variants with decreasing gPPA. We compared this to a random distribution of regulatory variants with respect to gPPA by permuting the CNN q-values 1000 times, preserving the overall number of significant variants, as well as the number and overall structure of credible sets. The enrichment p-value was calculated from these permutations by comparing the areas under curves with fraction of variants significant at each gPPA threshold generated in each permutation with the area under the curve calculated from the original results. Areas under curves were calculated using the AUC function of the DescTools R package (Signorell al., 2019). We repeated the same for functional fine-mapping PPAs (fPPA), using fPPAs generated through incorporation of pancreatic islet regulatory elements defined by chromatin state maps (Thurner et al., 2018). To account for the differing degree of fine-mapping resolution at different GWAS loci, we also evaluated whether we observed higher proportion of regulatory variants with q<0.05 among the variants with the highest PPA within each locus, regardless of the actual PPA value, and repeated the same procedure investigating genomic PPA ranks within each signal. In the same way we evaluated whether we observed lower q-values at the top ranks of signals known to act through insulin secretion mechanisms (N=34), rather than insulin action (N=23) (Dimas et al., 2014; Wood et al., 2017).

### Identification of T2D association signals further refined by incorporation of CNN regulatory prediction

We identified the T2D association signals where the islet CNN models can help with further refinement by investigating signals comprising at least 2 variants with fPPA>=0.2. In Figures 4 and 5 we highlighted signals where CNN predictions point to a single variant, among these fPPA>=0.2 variants, with a much higher regulatory score than the remaining variants at these loci.

The *in silico* saturated mutagenesis was performed using the basset_sat.py script from the Basset framework (Kelley et al., 2016). The TF binding motifs overlapping the variant site were identified with FIMO (Grant, Bailey, & Noble, 2011) using the CIS-BP motifs database (Weirauch et al., 2014).

### Plasmid transfection and Luciferase reporter assay

We experimentally validated the CNN regulatory predictions for the two variants (rs17712208 and rs79687284) at *PROX1* locus with luciferase reporter assay. Briefly, human EndoC-βH1 cells (Ravassard et al., 2011) were grown at 50–60% confluence in 24-well plates and were transfected with 500 ng of empty pGL4.23 [luc2/minP] vector (Promega, Charbonnieres, France) or pGL4-minP-PROX_enhancer vectors (wildtype, rs17712208-A and rs79687284-C) with FuGENE HD (Roche Applied Science, Meylan, France) using a FuGENE:DNA ratio of 6:1 according to manufacturer’s instruction, with three biological replicates for each tested condition. Primer sequences used for cloning and site-directed mutagenesis (SDM) are listed in STable 6. Restriction enzymes Nhel and Xhol were used for all subsequent cloning. Isolated clones were verified by sequencing.

Luciferase activities were measured 48 h after transfection using the Dual-Luciferase Reporter Assay kit (Promega) according to manufacturer’s instructions in half-volume 96-well tray format on an Enspire Multimode Plate Reader (PerkinElmer) with three technical replicates for each well. The firefly luciferase activity was normalized to the Renilla luciferase activity obtained by cotransfection of 10 ng of the pGL4.74[hRluc/TK] Renilla Luciferase vector (Promega). Results were analysed with two-tailed independent samples *t*-tests with *p*-value<0.05 considered as significantly different between groups.

### Comparison of single-versus multi-tissue CNN predictions

Finally, we compared the predictions resulting from our single-tissue pancreatic islet CNN models with the predictions generated with another publicly available multi-tissue variant prioritization tool DeepSEA (Zhou & Troyanskaya, 2015). The functional significance scores for each variant (multi-tissue) and the q-value for chromatin effects in the ENCODE PanIslets primary pancreatic islets cells (single tissue) were generated by submitting VCF files of the credible set variants to the DeepSEA web server (date accessed: 9^th^ May, 2018).

## Data availability

The datasets analysed during the current study are available in the public repositories under accessions listed in STable 1.

## Code availability

Code used to generate the results of this study is available at https://github.com/agawes/islet_CNN

## Acknowledgments

MT was a Wellcome doctoral student. ALG is a Wellcome Senior Fellow in Basic Biomedical Science. MMcC is a Wellcome Investigator and an NIHR Senior Investigator. Relevant funding support for this work comes from Wellcome (090532, 106130, 098381, 203141, 212259, 095101, 200837, 099673/Z/12/Z), Medical Research Council (MR/L020149/1), European Union Horizon 2020 Programme (T2D Systems), NIDDK (U01-DK105535), NIH (U01-DK105535; U01-DK085545) and NIHR (NF-SI-0617-10090).

## Author contributions

AWA and MMcC conceptualized the project, AWA performed all the computational analyses, and wrote the manuscript, GY and FA performed the *in vitro* assays, VN generated the ATAC-seq datasets, MT and JT performed the functional fine-mapping analysis, AM performed the genetic fine-mapping analysis, ALG and MMcC jointly supervised this work. All authors provided critical feedback to the manuscript.

---

**Figure 1–figure supplement 1.**
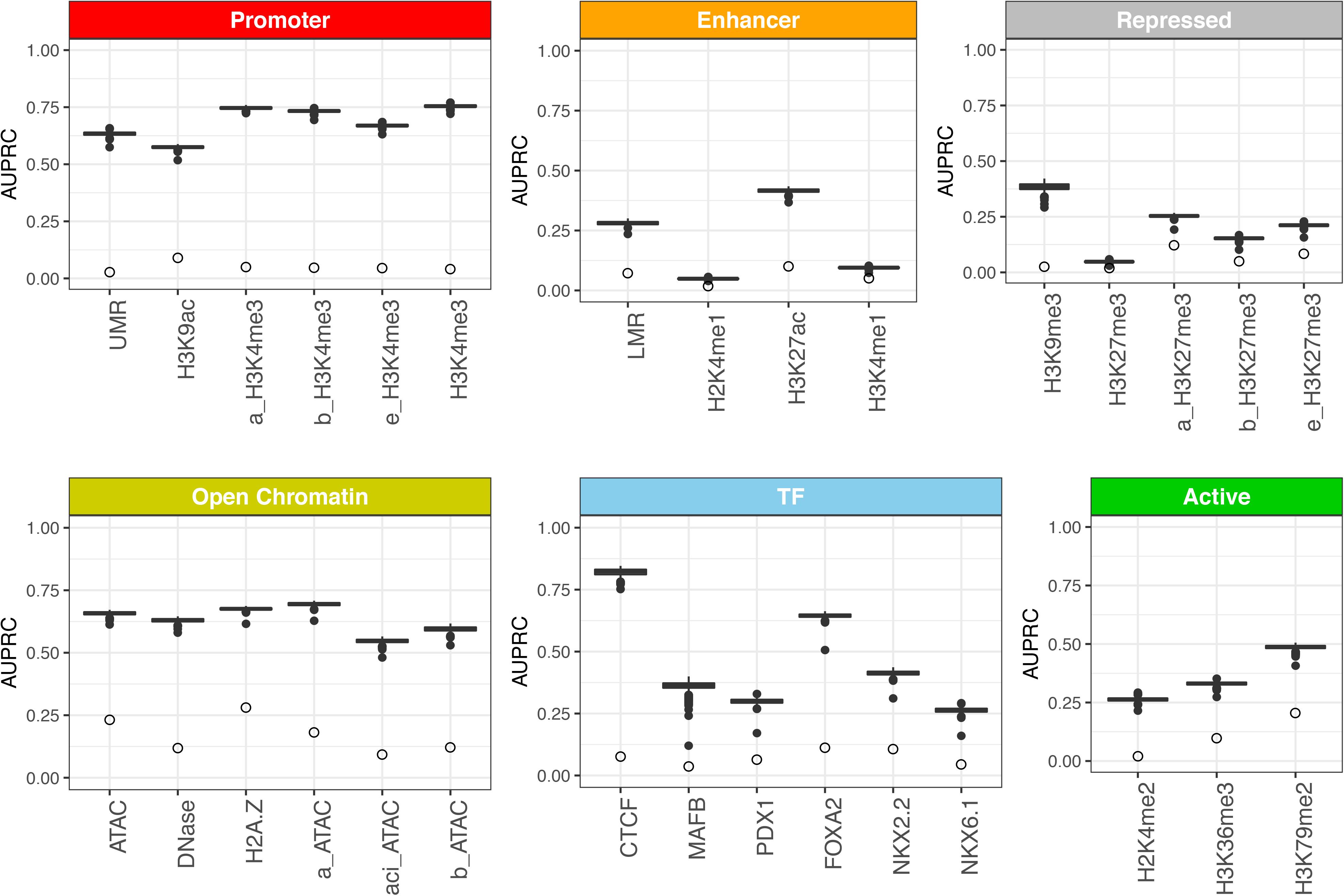
Area under precision-recall curves (AUPRC) for 30 islet epigenomic features predicted by CNN models. The AUPRC values were calculated based on performance on the validation set formed by 1000bp sequences from chr2, held out from training and testing. The boxplots show summary of performance across 1000 individual CNN models, and are grouped by corresponding regulatory element. As the interpretation of AUPRC values depend on how well balanced the dataset it, we denote the class imbalance (equivalent to prediction of a random model) for each feature as open circles.

**Figure 1–figure supplement 2.**
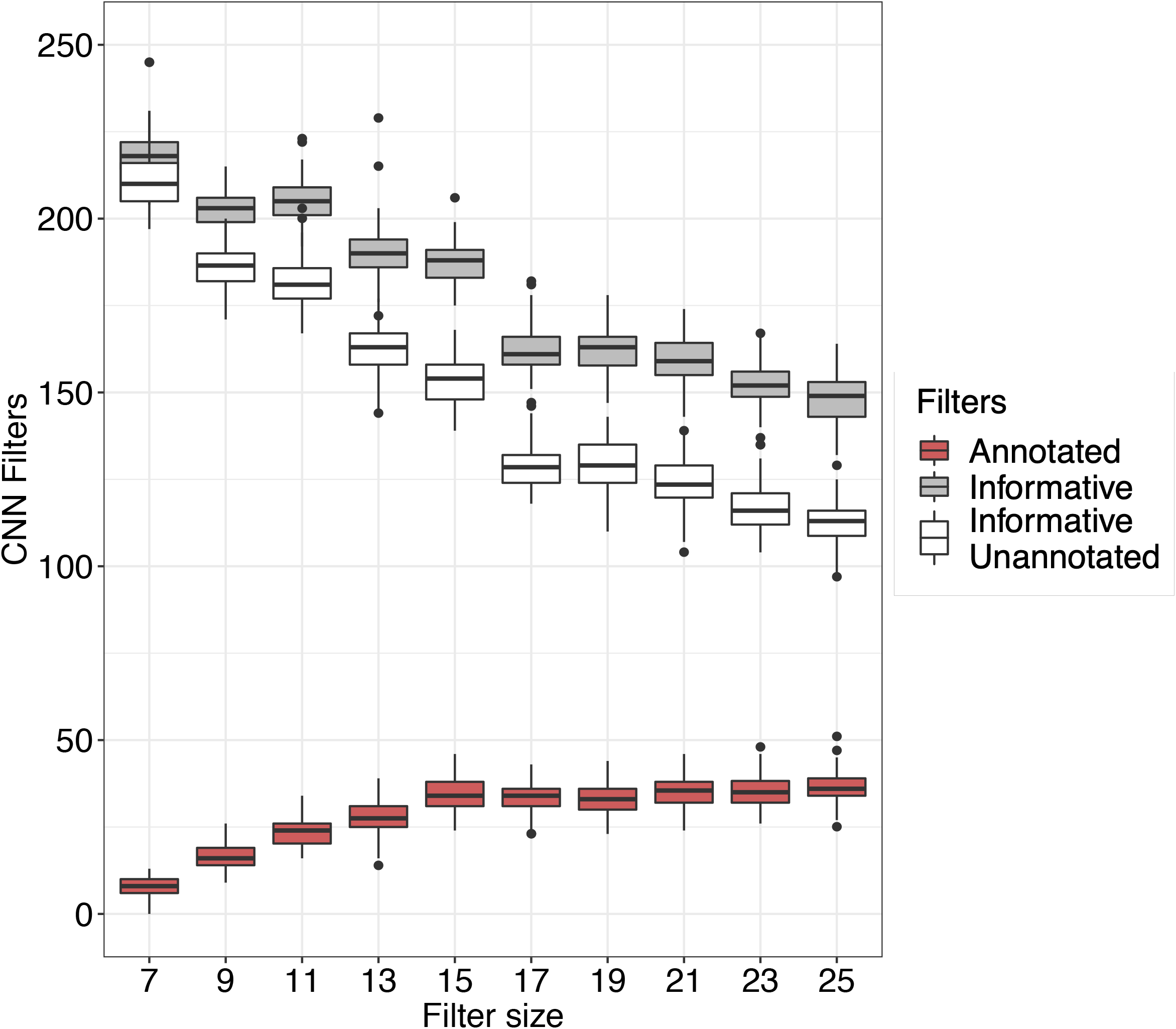
Influence of the size of filters in the first convolutional layers on filters’ annotation and filter’s influence on predictions. Boxplots represent summary of 100 individual CNN models differing in the size of convolutional filters of the first layer. In grey are shown all the informative filters, with standard deviation of filter activation >0, and in white a subset of these filters which were not annotated to match any known TF binding motifs. The number of informative filters decreases with increasing filter size. Red boxplots show increasing number of annotated filters with increasing filter size.

**Figure 3–figure supplement 1.**
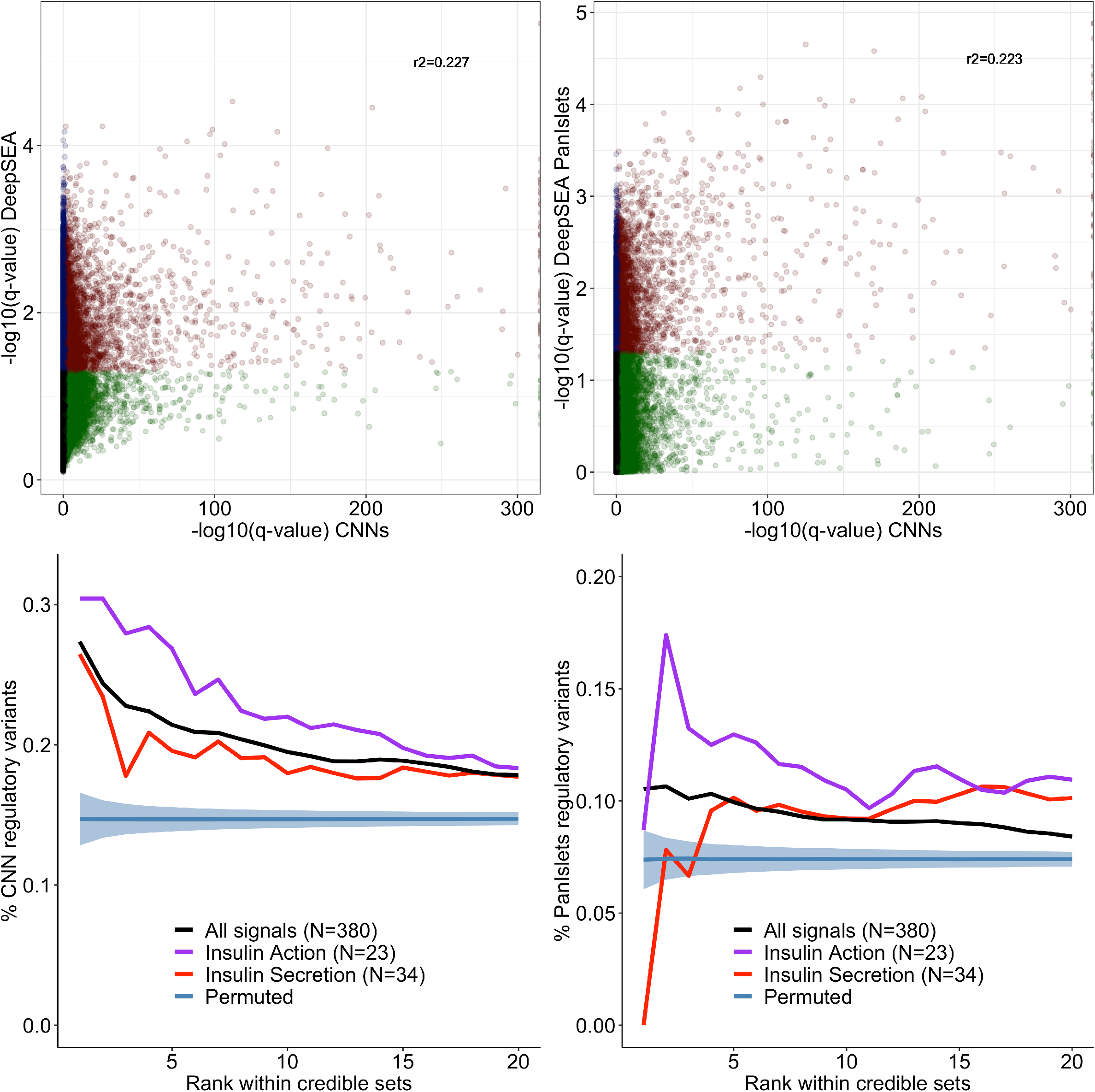
Comparison of CNN regulatory predictions made with the islet-specific CNN ensemble to predictions made with the publicly available DeepSEA model. A) Comparison of −log10-transformed q-values from the islet CNN ensemble with functional significance scores generated by the omni-tissue DeepSEA model B) Comparison of −log10-transformed q-values from the islet CNN ensemble with −log10-transformed E-values for the ENCODE PanIslet DNase generated by the DeepSEA model. In scatterplots variants predicted to be regulatory with both approaches are shown in red, variants predicted as regulatory only by DeepSEA are shown in blue, and variants predicted as regulatory only by islet CNNs are shown in green. C) Enrichment of regulatory variants among variants at the top ranks of T2D GWAS credible sets (black) predicted by the omni-tissue DeepSEA model over the permuted background (blue). Purple line shows the signals acting through insulin action mechanisms, while red line shows the signals acting through insulin secretion (pancreatic islet-mediated) mechanisms. D) Enrichment of regulatory variants among variants at the top ranks of T2D GWAS credible sets (black) predicted by the single-tissue DeepSEA model based on ENCODE PanIslet DNase dataset over the permuted background (blue). Purple line shows the signals acting through insulin action mechanisms, while red line shows the signals acting through insulin secretion (pancreatic islet-mediated) mechanisms.

## References

Ackermann, A. M., Wang, Z., Schug, J., Naji, A., & Kaestner, K. H. (2016). Integration of ATAC-seq and RNA-seq identifies human alpha cell and beta cell signature genes. Mol Metab, 5(3), 233–244. doi:10.1016/j.molmet.2016.01.002

Bernstein, B. E., Stamatoyannopoulos, J. A., Costello, J. F., Ren, B., Milosavljevic, A., Meissner, A., … Thomson, J. A. (2010). The NIH Roadmap Epigenomics Mapping Consortium. Nat Biotechnol, 28(10), 1045–1048. doi:10.1038/nbt1010-1045

Bhandare, R., Schug, J., Le Lay, J., Fox, A., Smirnova, O., Liu, C., … Kaestner, K. H. (2010). Genome-wide analysis of histone modifications in human pancreatic islets. Genome Res, 20(4), 428–433. doi:10.1101/gr.102038.109

Bramswig, N. C., Everett, L. J., Schug, J., Dorrell, C., Liu, C., Luo, Y., … Kaestner, K. H. (2013). Epigenomic plasticity enables human pancreatic alpha to beta cell reprogramming. J Clin Invest, 123(3), 1275–1284. doi:10.1172/JCI66514

Cooper, G. M., Stone, E. A., Asimenos, G., Program, N. C. S., Green, E. D., Batzoglou, S., & Sidow, A. (2005). Distribution and intensity of constraint in mammalian genomic sequence. Genome Res, 15(7), 901–913. doi:10.1101/gr.3577405

Dimas, A. S., Lagou, V., Barker, A., Knowles, J. W., Magi, R., Hivert, M. F., … Investigators, M. (2014). Impact of type 2 diabetes susceptibility variants on quantitative glycemic traits reveals mechanistic heterogeneity. Diabetes, 63(6), 2158–2171. doi:10.2337/db13-0949

Gaulton, K. J., Ferreira, T., Lee, Y., Raimondo, A., Magi, R., Reschen, M. E., … Meta-analysis, C. (2015). Genetic fine mapping and genomic annotation defines causal mechanisms at type 2 diabetes susceptibility loci. Nat Genet, 47(12), 1415–1425. doi:10.1038/ng.3437

Gaulton, K. J., Nammo, T., Pasquali, L., Simon, J. M., Giresi, P. G., Fogarty, M. P., … Ferrer, J. (2010). A map of open chromatin in human pancreatic islets. Nat Genet, 42(3), 255–259. doi:10.1038/ng.530

Grant, C. E., Bailey, T. L., & Noble, W. S. (2011). FIMO: scanning for occurrences of a given motif. Bioinformatics, 27(7), 1017–1018. doi:10.1093/bioinformatics/btr064

Grau, J., Grosse, I., & Keilwagen, J. (2015). PRROC: computing and visualizing precision-recall and receiver operating characteristic curves in R. Bioinformatics, 31(15), 2595–2597. doi:10.1093/bioinformatics/btv153

GTex Consortium, Laboratory, Data Analysis & Coordinating Center-Analysis Working groups, Statistical Methods groups-Analysis Working Group, Enhancing GTex (eGTex) groups, NIH Common Fund, … Montgomery, S. B. (2017). Genetic effects on gene expression across human tissues. Nature, 550(7675), 204–213. doi:10.1038/nature24277

Gu, C., Stein, G. H., Pan, N., Goebbels, S., Hornberg, H., Nave, K. A., … Lee, J. E. (2010). Pancreatic beta cells require NeuroD to achieve and maintain functional maturity. Cell Metab, 11(4), 298–310. doi:10.1016/j.cmet.2010.03.006

Gupta, S., Stamatoyannopoulos, J. A., Bailey, T. L., & Noble, W. S. (2007). Quantifying similarity between motifs. Genome Biol, 8(2), R24. doi:10.1186/gb-2007-8-2-r24

Huang, C., Thompson, P., Wang, Y., Yu, Y., Zhang, J., Kong, D., … Alzheimer’s Disease Neuroimaging, I. (2017). FGWAS: Functional genome wide association analysis. Neuroimage, 159, 107–121. doi:10.1016/j.neuroimage.2017.07.030

Jennings, R. E., Berry, A. A., Strutt, J. P., Gerrard, D. T., & Hanley, N. A. (2015). Human pancreas development. Development, 142(18), 3126–3137. doi:10.1242/dev.120063

Kelley, D. R., Reshef, Y. A., Bileschi, M., Belanger, D., McLean, C. Y., & Snoek, J. (2018). Sequential regulatory activity prediction across chromosomes with convolutional neural networks. Genome Res, 28(5), 739–750. doi:10.1101/gr.227819.117

Kelley, D. R., Snoek, J., & Rinn, J. L. (2016). Basset: learning the regulatory code of the accessible genome with deep convolutional neural networks. Genome Res, 26(7), 990–999. doi:10.1101/gr.200535.115

Liu, L., Sanderford, M. D., Patel, R., Chandrashekar, P., Gibson, G., & Kumar, S. (2019). Biological relevance of computationally predicted pathogenicity of noncoding variants. Nat Commun, 10(1), 330. doi:10.1038/s41467-018-08270-y

Luo, W., Friedman, M. S., Shedden, K., Hankenson, K. D., & Woolf, P. J. (2009). GAGE: generally applicable gene set enrichment for pathway analysis. BMC Bioinformatics, 10, 161. doi:10.1186/1471-2105-10-161

Mahajan, A., Taliun, D., Thurner, M., Robertson, N. R., Torres, J. M., Rayner, N. W., … McCarthy, M. I. (2018). Fine-mapping type 2 diabetes loci to single-variant resolution using high-density imputation and islet-specific epigenome maps. Nat Genet, 50(11), 1505–1513. doi:10.1038/s41588-018-0241-6

Maher, B. (2012). ENCODE: The human encyclopaedia. Nature, 489(7414), 46–48.

Marbach, D., Lamparter, D., Quon, G., Kellis, M., Kutalik, Z., & Bergmann, S. (2016). Tissue-specific regulatory circuits reveal variable modular perturbations across complex diseases. Nat Methods, 13(4), 366–370. doi:10.1038/nmeth.3799

Miguel-Escalada, I., Bonàs-Guarch S., Cebola I., Ponsa-Cobas J., Mendieta-Esteban J., Rolando D.M.Y, Javierre B.J., Atla G., Farabella I., Morgan C.C., García-Hurtado J., Beucher A., Morán I., Pasquali L., Ramos M., Appel E.V.R., Linneberg A., Gjesing A.P., Witte D.R., Pedersen O., Garup N., Ravassard P., Torrents D., Mercader J.M., Piemonti L., Berney T., de Koning E.J.P., Kerr-Conte J., Pattou F., Fedko I.O., Prokopenko I., Hansen T., Marti-Renom M.A., Fraser P., Ferrer J. (2018). Human pancreatic islet 3D chromatin architecture provides insights into the genetics of type 2 diabetes. bioRxiv. doi:https://doi.org/10.1101/400291

Parker, S. C., Stitzel, M. L., Taylor, D. L., Orozco, J. M., Erdos, M. R., Akiyama, J. A., … Authors, N. C. S. P. (2013). Chromatin stretch enhancer states drive cell-specific gene regulation and harbor human disease risk variants. Proc Natl Acad Sci U S A, 110(44), 17921–17926. doi:10.1073/pnas.1317023110

Pasquali, L., Gaulton, K. J., Rodriguez-Segui, S. A., Mularoni, L., Miguel-Escalada, I., Akerman, I., … Ferrer, J. (2014). Pancreatic islet enhancer clusters enriched in type 2 diabetes risk-associated variants. Nat Genet, 46(2), 136–143. doi:10.1038/ng.2870

Ravassard, P., Hazhouz, Y., Pechberty, S., Bricout-Neveu, E., Armanet, M., Czernichow, P., & Scharfmann, R. (2011). A genetically engineered human pancreatic beta cell line exhibiting glucose-inducible insulin secretion. J Clin Invest, 121(9), 3589–3597. doi:10.1172/JCI58447

Signorell, A. S. e. et al. (2019). DescTools: Tools fro descriptive statistics. R package version 0.99.28. Retrieved from https://cran.r-project.org/package=DescTools

Stitzel, M. L., Sethupathy, P., Pearson, D. S., Chines, P. S., Song, L., Erdos, M. R., … Collins, F. S. (2010). Global epigenomic analysis of primary human pancreatic islets provides insights into type 2 diabetes susceptibility loci. Cell Metab, 12(5), 443–455. doi:10.1016/j.cmet.2010.09.012

Tewhey, R., Kotliar, D., Park, D. S., Liu, B., Winnicki, S., Reilly, S. K., … Sabeti, P. C. (2016). Direct Identification of Hundreds of Expression-Modulating Variants using a Multiplexed Reporter Assay. Cell, 165(6), 1519–1529. doi:10.1016/j.cell.2016.04.027

Thurner, M., van de Bunt, M., Torres, J. M., Mahajan, A., Nylander, V., Bennett, A. J., … McCarthy, M. I. (2018). Integration of human pancreatic islet genomic data refines regulatory mechanisms at Type 2 Diabetes susceptibility loci. Elife, 7. doi:10.7554/eLife.31977

Ulirsch, J. C., Nandakumar, S. K., Wang, L., Giani, F. C., Zhang, X., Rogov, P., … Sankaran, V. G. (2016). Systematic Functional Dissection of Common Genetic Variation Affecting Red Blood Cell Traits. Cell, 165(6), 1530–1545. doi:10.1016/j.cell.2016.04.048

van der Meulen, T., & Huising, M. O. (2015). Role of transcription factors in the transdifferentiation of pancreatic islet cells. J Mol Endocrinol, 54(2), R103–117. doi:10.1530/JME-14-0290

Weirauch, M. T., Yang, A., Albu, M., Cote, A. G., Montenegro-Montero, A., Drewe, P., … Hughes, T. R. (2014). Determination and inference of eukaryotic transcription factor sequence specificity. Cell, 158(6), 1431–1443. doi:10.1016/j.cell.2014.08.009

Wood, A. R., Jonsson, A., Jackson, A. U., Wang, N., van Leewen, N., Palmer, N. D., … Frayling, T. M. (2017). A Genome-Wide Association Study of IVGTT-Based Measures of First-Phase Insulin Secretion Refines the Underlying Physiology of Type 2 Diabetes Variants. Diabetes, 66(8), 2296–2309. doi:10.2337/db16-1452

Zhou, J., Theesfeld, C. L., Yao, K., Chen, K. M., Wong, A. K., & Troyanskaya, O. G. (2018). Deep learning sequence-based ab initio prediction of variant effects on expression and disease risk. Nat Genet, 50(8), 1171–1179. doi:10.1038/s41588-018-0160-6

Zhou, J., & Troyanskaya, O. G. (2015). Predicting effects of noncoding variants with deep learning-based sequence model. Nat Methods, 12(10), 931–934. doi:10.1038/nmeth.3547

